# Synergistic interactions between *Stachybotrys chartarum* and other indoor fungal species in moisture-damaged houses

**DOI:** 10.1101/2021.07.13.452289

**Authors:** Paris Chakravarty

## Abstract

The interactions between six commonly occurring fungal species in damp or water-damaged houses in southern California were studied. These fungal species were *Alternaria alternata, Aspergillus niger, Chaetomium globosum, Cladosporium herbarum, Penicillium chrysogenum* and *Stachybotrys chartarum.* In the damp building materials, *S. chartarum* was found to be associated with *A. niger, C. globosum,* and *P. chrysogenum* but not with *A. alternata* and *C. herbarum*. *Stachybotrys chartarum* showed strong antagonistic effect against *A. alternata* and *C. herbarum* and significantly inhibited *in vitro* growth of *A. alternata* and *C. herbarum* but had no effect on *A. niger, C. globosum,* and *P. chrysogenum*. Two trichothecenes, produced by *S. chartarum*, trichodermin and trichodermol, significantly inhibited spore germination and *in vitro* growth of *A. alternata* and *C. herbarum* but had no effect on *A. niger, C. globosum, P. chrysogenum* and *S. chartarum.* In the damp building materials (drywall, ceiling tile, and oak woods), *S. chartarum* significantly inhibited the growth of *A. alternata* and *C. herbarum* and had no effect on the growth and colonization of *A. niger, C. globosum, P. chrysogenum* in these substrata.

## INTRODUCTION

Fungal exposures in damp or water-damaged houses have become a major concern because of their potential health effects and there is clear relationship between contaminated indoor environments and illness. Indoor air quality became an important issue since 1960s when researchers found that indoor pollutant levels in damp houses can reach or exceed those of outdoor levels (28). It is now well known fact that indoor air quality is an essential part of our health since we spend 90% of our time indoors inhaling approximately 15 m^3^ of ambient air every day (20). It is reported that the occupants of wet, moldy buildings have increase in subjective complaints and children in damp homes show higher respiratory and other illness (4,17,23). The list of symptoms generally consists of upper respiratory complaints, including headache, eye irritation, epistaxis, nasal and sinus congestion, cough, and gastrointestinal complaints (19). The exposure to fungal spores enhances the histamine release triggered by both allergic and non-immunologic mechanisms in the cultured leukocytes (7). Besides moisture damage, high temperature and relative humidity can also contribute to the higher occurrence of indoor fungi in the house (5,25).

During our indoor air quality study at the moisture-damaged houses in southern California, we have found six commonly occurring fungal species in the indoor air and building materials were *Alternaria alternata* (Fr.) Keissl., *Aspergillus niger* van Tieghem, *Chaetomium globosum* Kunze ex Steud., *Cladosporium herbarum* (Pers.) Link ex Gray, *Penicillium chrysogenum* Thom, and *Stachybotrys chartarum* (Ehrenb. ex Link) Hughes (6). It was found that certain fungi occur alone or in combination with other fungi. Interactions between the different fungi in a moisture-damaged house are inevitable because spores of a single fungal species alone may contain various metabolites, and moisture-damaged site is always a habitat of more than one fungal species (3,6,21). Many mycotoxins are thought to be involved in chemical signaling between organisms or species and the production of some of the mycotoxins may be stimulated or inhibited when microorganisms are interact with each other (8, 22). We have found that *S. chartarum* was growing alone or in combination with *A. niger, C. globosum,* and *P. chrysogenum* in the damp or water-damaged building materials but not with *A. alternata* and *C. herbarum.* The frequent occurrence of *S. chartarum* with *A. niger, C. globosum,* and *P. chrysogenum* in moisture-damaged building materials means that they often share their habitat in common damp or water-damaged building materials without any effect on the mycotoxin produced by *S. chartarum*. *Stachybotrys chartarum* produce secondary metabolites known as trichothecenes that are harmful to human and animal health and affects at cellular level (1,2,9,18,27). Several compounds in trichothecenes family have been isolated including trichodermin and trichodermol (15,16) which are precursors to the satratoxin, a highly cytotoxic compound that suppresses the immune system.

The objective of this study was to investigate the synergistic interactions between *S. chartarum* and its secondary metabolites trichodermin and trichodermol on five commonly occurring fungal species in damp building materials colonized by *A. alternata, A. niger, C. globosum, C. herbarum*, and *P. chrysogenum* on their growth, spore germination, and their ability to grow and multiply in the damp or water-damaged building materials.

## MATERIALS AND METHODS

### Fungal species and mycotoxin

Six commonly occurring fungal species found in the moisture-damaged houses in southern California were used in this study. These fungal species were *A. alternata, A. niger, C. globosum, C. herbarum, P. chrysogenum* and *S. chartarum.* Mycotoxins used in this study were trichodermin and trichodermol belong to trichothecene family produced by *S. chartarum*.

### *In vitro* antagonism

Antagonism of *S. chartarum* against *A. alternata, A. niger, C. globosum, C. herbarum*, and *P. chrysogenum* was studied on malt extract agar (MEA), potato dextrose agar (PDA) and carrot agar (CA) media in 90-mm Petri plates. *Stachybotrys chartarum* (a slower growing fungus) was inoculated with 5-mm agar plugs at the margin of the plate and allowed to grow at 25° C in the dark. For each medium, there were 20 replicates. Seven days later, 5-mm mycelial disks of *A. alternata, A. niger, C. globosum, C. herbarum*, and *P. chrysogenum* from agar cultures were placed separately on the agar plates opposite to *S. chartarum* and the Petri plates were then incubated at described above. Colony diameter of the fungi was measured 7 days later using a ruler. The inhibition zone formed around *A. alternata, A. niger, C. globosum, C. herbarum*, and *P. chrysogenum* were measured in a straight line from the edge of the *S. chartarum* colony to the edge of the *A. alternata, A. niger, C. globosum, C. herbarum*, and *P. chrysogenum* colonies 7 days later.

### Effect of culture filtrates of *S. chartarum* on the *in vitro* growth of five species of fungi in agar diffusion plates

The effect of culture filtrates of *S. chartarum* on *A. alternata, A. niger, C. globosum, C. herbarum*, and *P. chrysogenum* was studied on agar diffusion plates. These plates were prepared from 90 mm Petri plates containing 40 ml of MEA, PDA, and CA media, by removing 5 mm diameter agar plugs from each of four quarters of the plate. Five mm diameter agar plugs of *A. alternata, A. niger, C. globosum, C. herbarum*, and *P. chrysogenum* were separately inoculated in the center of the agar diffusion plates, and incubated at 25° C in the dark. The culture filtrate of *S. chartarum* was prepared by filtering 15-day-old liquid cultures (grown in carrot extract (CE) in 250 ml flasks on a shaker) through both Whatman No. 1 filter paper and a 0.45 μm Milipore filter and then drying on a rotary evaporator at 45° C. The evaporated sample was resuspended in 5 ml distilled water. One ml of filter sterilized concentrated culture filtrate of *S. chartarum* was added to diffusion wells of each of the 20 replicate plates containing 3-day old cultures of *A. alternata, A. niger, C. globosum, C. herbarum*, and *P. chrysogenum.* The plates were incubated at 25° C in the dark. After incubation for 7 days, the zone of inhibition around each diffusion well was measured. Microscopic examination of hyphae of *A. alternata, A. niger, C. globosum, C. herbarum*, and *P. chrysogenum* were also made.

### Effect of culture filtrates of *S. chartarum* on the *in vitro* mycelial growth of five species of fungi

For this experiment, *S. chartarum* was grown in liquid malt extract (ME), potato dextrose broth (PDB), and CE at 25° C in the dark. After 15 days, the mycelia were harvested on Whatman No. 1 filter paper. The culture filtrate was collected and stored at 2° C in the dark overnight. Fifty ml of liquid media (ME, PDB, and CE) were autoclaved in 250-ml flasks for 15 min at 121° C. When cooled down, flasks were separately inoculated with 5 agar plugs (5 mm diameter) of actively growing mycelia of *A. alternata, A. niger, C. globosum, C. herbarum*, and *P. chrysogenum.* The agar was removed with a sterile scalpel and only mycelial mats were inoculated in the flasks. Five ml of culture filtrate of *S. chartarum* was then added to each flask. The control consisted of 5 ml of sterile distilled water. The flasks were kept in the dark on a shaker at room temperature (24 ± 2° C). After 4-week incubation period, the mycelia of *A. alternata, A. niger, C. globosum, C. herbarum*, and *P. chrysogenum* were harvested on a Whatmen No 1 filter paper, oven dried at 70° C for 48 hr and the dry weight of mycelia calculated.

### Effect of trichodermin and trichdermol on the spore germination and *in vitro* growth of six species of fungi

To test the effect of trichodermin and trichodermol on the spore germination, *A. alternata, A. niger, C. globosum, C. herbarum*, and *P. chrysogenum* were grown on 2% MEA at 25° C in the dark for 1 week, whereas *S. chartarum* was grown on CA for 2 weeks. The spore suspension was prepared by transferring from fungal culture with a transfer loop into 9 ml sterile distilled water and the concentration of the suspension was adjusted to approximately 10^5^ spores/ml. Ten μl of spore suspension of *A. alternata, A. niger, C. globosum, C. herbarum, P. chrysogenum* and *S. chartarum* was mixed separately with ten μl of filter sterilized trichodermin and trichodermol in a cavity slide. The slides with spores were kept moist by placing them on glass rods on the moistened filter paper in Petri plates and sealed with parafilm. Spore germination was recorded after 24 hr incubation at 25° C in the dark, and 100 spores were counted for each treatment.

To test the effect of trichodermin and trichodermol on the *in vitro* growth of *A. alternata, A. niger, C. globosum, C. herbarum, P. chrysogenum*, and *Stachybotrys,* these fungi were grown on multiwall tissue culture plates (1.8 × 1.5 cm diameter X length of individual wells). Two per cent MEA was used for *A. alternata, A. niger, C. globosum, C. herbarum*, and *P. chrysogenum* and for *S. chartarum*, CA was used. Twenty five μl of trichodermin and trichodermol at 1, 10, 100, and 1000 ppm in acetone-dichloromethane was added separately to the surface of the nutrient media in each well. For each concentration, 15 multiwells were used. In the control, only 25 μl of acetone-dichloromethane was used. All the multiwells were kept in a laminar flow hood for 1 min to allow the solvents to evaporate. Each agar well was individually inoculated with 5-mm agar plugs of *A. alternata, A. niger, C. globosum, C. herbarum, P. chrysogenum* and *S. chartarum* and then multiwells were wrapped with parafilm and incubated at 25° C in the dark. After 5 days of incubation, the colony diameter of *A. alternata, A. niger*, *C. globosum, C. herbarum*, and *P. chrysogenum* were measured and mycelia were observed under a microscope. For *S. chartarum* colony diameter and microscopic observation were made after 10 days.

### Effect of *S. chartarum* towards five fungal species on colonization in the building materials

#### Growth of fungal species in the building materials

Three building materials (powdered drywall, powdered ceiling tile, and oak wood chips) were used in this study. The materials were crushed into coarse material using a grinder. One hundred gm of drywall, ceiling tile, and wood chips were soaked separately in ME in each of 95 flasks. After 1 hr, the ME was drained and flasks were autoclaved for 60 min at 121° C. There were 5 replicates for each treatment. When cooled, 30 flasks containing dry wall, 30 flasks containing ceiling tile, and 30 flasks containing oak wood chips were separately inoculated with 5 agar plugs (5-mm diameter) of actively growing *A. alternata, A. niger, C. globosum, C. herbarum, P. chrysogenum* and *S*. *chartarum*. Five control flasks received 5 agar plugs (5-mm diameter) without any fungal species growing on it. The flasks were kept in the incubator at 25° C in the dark. After 30 days of incubation, flasks were removed from the incubator, observed under a microscope, and isolation of the fungi was made from the inoculated drywall, ceiling tile, and wood chips.

#### Interaction of S. chartarum on the growth of five fungal species in the building materials

Three building materials described above were used in this study. One hundred gm of drywall, ceiling tile, and wood chips were soaked separately in ME in each eighty 250 ml flasks. After 1 hr, the ME was drained and flasks were autoclaved for 60 min at 121° C. There were 5 replicates for each treatment. When cooled, 25 flasks containing dry wall, 25 flasks containing ceiling tile, and 25 flasks containing oak wood chips were aseptically inoculated with 5 agar plugs (5-mm diameter) of actively growing mycelia of *S. chartarum*. The flasks were incubated at 25° C in the dark. The flasks were shaken periodically to fragment the mycelia of the fungi on the drywall, ceiling tile, and wood chips. After 28 days, 5 flasks containing each of *S*. *chartarum* growing on drywall, ceiling tile and wood chips were separately inoculated with five agar plugs (5 mm diameter) of *A. alternata, A. niger, C. globosum, C. herbarum*, and *P. chrysogenum* and returned to the incubator. Five control flasks contained only *S. chartarum* did not receive any treatment. The following treatments resulted: *S*. *chartarum* + *A. alternata, S*. *chartarum* + *A. niger, S*. *chartarum* + *C. globosum, S*. *chartarum* + *C. herbarum, S*. *chartarum* + *P. chrysogenum* and only *S. chartarum*. After 30 days of incubation, flasks were removed from the incubator, observed under a microscope, and isolation of the fungi was made from the inoculated drywall, ceiling tile, and wood chips.

### Statistical analysis

Data were subjected to analysis of variance (31). Individual means were compared using Scheffe’s test for multiple comparisons using SAS software, version 9.0 (24). Means followed by the same letters (a,b,c and so on) in tables and bars (in the graphs) for a particular fungal species against *S. chartarum* and its metabolite trichodermin and trichodermol are not significantly (P=0.05) different from each other by Scheffe’s test for multiple comparison.

## RESULTS

### *In vitro* antagonism

#### Inhibitory effect of *Stachybotrys chartarum* towards five fungal species

The *in vitro* growth of *A. alternata* and *C. herbarum* was significantly inhibited when grown in dual culture with *S. chartarum* on all three culture media used (Table 1). The inhibition zones, formed around the colonies of the fungi ranged from 9.5 mm to 23.5 mm for *A. alternata* and 8.0 mm to 25.5 mm for *C. herbarum* in all three nutrient media tested (Table 1). *Aspergillus niger, C. globosum,* and *P. chrysogenum* had no effect on their growth when grown together with or without *S. chartarum* and no inhibition zones were formed around the colonies of *A. niger, C. globosum,* and *P. chrysogenum* (Table 1).

**TABLE 1.** Inhibitory effect of *Stachybotrys chartarum* towards five fungal species on different nutrient media in agar plates

| Fungal species | Nutrient Media | Dual culture with *S. chartarum* |  | Without *S. chartarum* |
| --- | --- | --- | --- | --- |
|  |  | Colony diameter (mm) | Inhibition zone (mm) | Colony diameter (mm) |
| *A. alternata* | MEA | 51.0b | 9.5c | 69.5a |
|  | PDA | 53.5b | 17.0c | 70.5a |
|  | CA | 50.5b | 23.5c | 60.0a |
| *A. niger* | MEA | 90.0a | 0.0b | 90.0a |
|  | PDA | 89.0a | 0.0b | 88.0a |
|  | CA | 87.5a | 0.0b | 88.5a |
| *C. globosum* | MEA | 40.5a | 0.0b | 42.0a |
|  | PDA | 36.0b | 0.0c | 41.5a |
|  | CA | 33.5b | 0.0c | 39.5a |
| *C. herbarum* | MEA | 44.0b | 8.0c | 63.0a |
|  | PDA | 42.0b | 21.0c | 54.0a |
|  | CA | 36.5b | 25.5c | 46.5a |
| *P. chrysogenum* | MEA | 89.5a | 0.0b | 90.0a |
|  | PDA | 90.0a | 0.0b | 89.0a |
|  | CA | 88.5a | 0.0b | 90.0a |

Values are the means of 20 replicates. Means followed by the same letters (a,b,c) in a row for a particular species are not significantly ( $P=0.05$ ) different from each other by Scheffe's test for multiple comparison. MEA, malt extract agar; PDA, potato dextrose agar; CA, carrot agar.

Similarly, in agar diffusion plates, inhibition zone was observed when *A. alternata* and *C. herbarum* were treated with culture filtrate of *S. chartarum* (Table 2). For *A. alternata* inhibition zone ranged from 6.5 mm to 25.5 mm and for *C. herbarum* it was 15.0 mm to 31.5 mm (Table 2). No inhibition zone was observed for *A. niger, C. globosum,* and *P. chrysogenum* (Table 2). Control plates without culture filtrate of *S. chartarum* had no inhibition zones (Table 2).

**TABLE 2.** Inhibitory effect of culture filtrate of *Stachybotrys chartarum* towards five fungal species on different nutrient media in agar diffusion plates

| Fungal species | Nutrient Media | Culture Filtrate of *S. chartarum*<br>Inhibition zone (mm) | Control (Without Culture Filtrate of *S. chartarum*)<br>Inhibition zone (mm) |
| --- | --- | --- | --- |
| *A. alternata* | MEA | 6.5b | 0.0a |
|  | PDA | 18.0b | 0.0a |
|  | CA | 25.5b | 0.0a |
| *A. niger* | MEA | 0.0a | 0.0a |
|  | PDA | 0.0a | 0.0a |
|  | CA | 0.0a | 0.0a |
| *C. globosum* | MEA | 0.0a | 0.0a |
|  | PDA | 0.0a | 0.0a |
|  | CA | 0.0a | 0.0a |
| *C. herbarum* | MEA | 15.0b | 0.0a |
|  | PDA | 28.0b | 0.0a |
|  | CA | 31.5b | 0.0a |
| *P. chrysogenum* | MEA | 0.0a | 0.0a |
|  | PDA | 0.0a | 0.0a |
|  | CA | 0.0a | 0.0a |

#### Effect of culture filtrates of *S. chartarum* on the *in vitro* mycelial growth of five species of fungi

The mycelial growth of *A. alternata* and *C. herbarum* was significantly inhibited when treated with culture filtrate of *S. chartarum* (Table 3). Vacuolation of the hyphae of *A. alternata* and *C. herbarum* was observed under a microscope. The mycelial growth of *A. niger, C. globosum,* and *P. chrysogenum* was not affected when grown with or without culture filtrate of *S. chartarum*. (Table 3). No vacuolation of the hyphae of *A. niger, C. globosum,* and *P. chrysogenum* was observed under a microscope when grown together with culture filtrate of *S. chartarum*.

**TABLE 3.** Effect of culture filtrate of *Stachybotrys chartarum* on the *in vitro* mycelial growth (mg dry wt) of five fungal species on different nutrient media

| Fungal species | Without *S. chartarum*<br>(mg mycelial dry wt.) |  |  | With *S. chartarum*<br>(mg mycelial dry wt.) |  |  |
| --- | --- | --- | --- | --- | --- | --- |
|  | ME | PDB | CE | ME | PDB | CE |
| *A. alternata* | 5.0b | 9.0a | 6.5b | 1.0c | 1.5c | 1.0c |
| *A. niger* | 103.0a | 96.0b | 87.5c | 104.0a | 98.0b | 86.5c |
| *C. globosum* | 21.0c | 30.5a | 27.0b | 22.5c | 29.0a | 25.5b |
| *C. herbarum* | 19.5a | 18.0a | 19.0a | 1.5b | 1.0b | 1.0b |
| *P. chrysogenum* | 86.0b | 91.0a | 74.0c | 85.0b | 89.5a | 73.0c |

Values are the means of 20 replicates. Means followed by the same letters (a,b,c) in a row for a particular species are not significantly ( $P=0.05$ ) different from each other by Scheffe's test for multiple comparison. ME, malt extract; PDB, potato dextrose broth; CE, carrot agar.

#### Effect of trichodermin and trichdermol on the spore germination and *in vitro* growth of five species of fungi

At 1 ppm of trichodermin and trichodermol, spore germination of *A. alternata* and *C. herbarum* was not affected (Figure 1 and Figure 2). Spore germination of *A. alternata* and *C. herbarum* was significantly reduced when treated with trichodermin and trichodermol at 10, 100, and 1000 ppm (Figure 1 and Figure 2). Both these compounds had no effect on spore germination of *A. niger, C. globosum, P. chrysogenum* and *S. chartarum* at 1, 10, 100, and 1000 ppm.

**FIG. 1.**
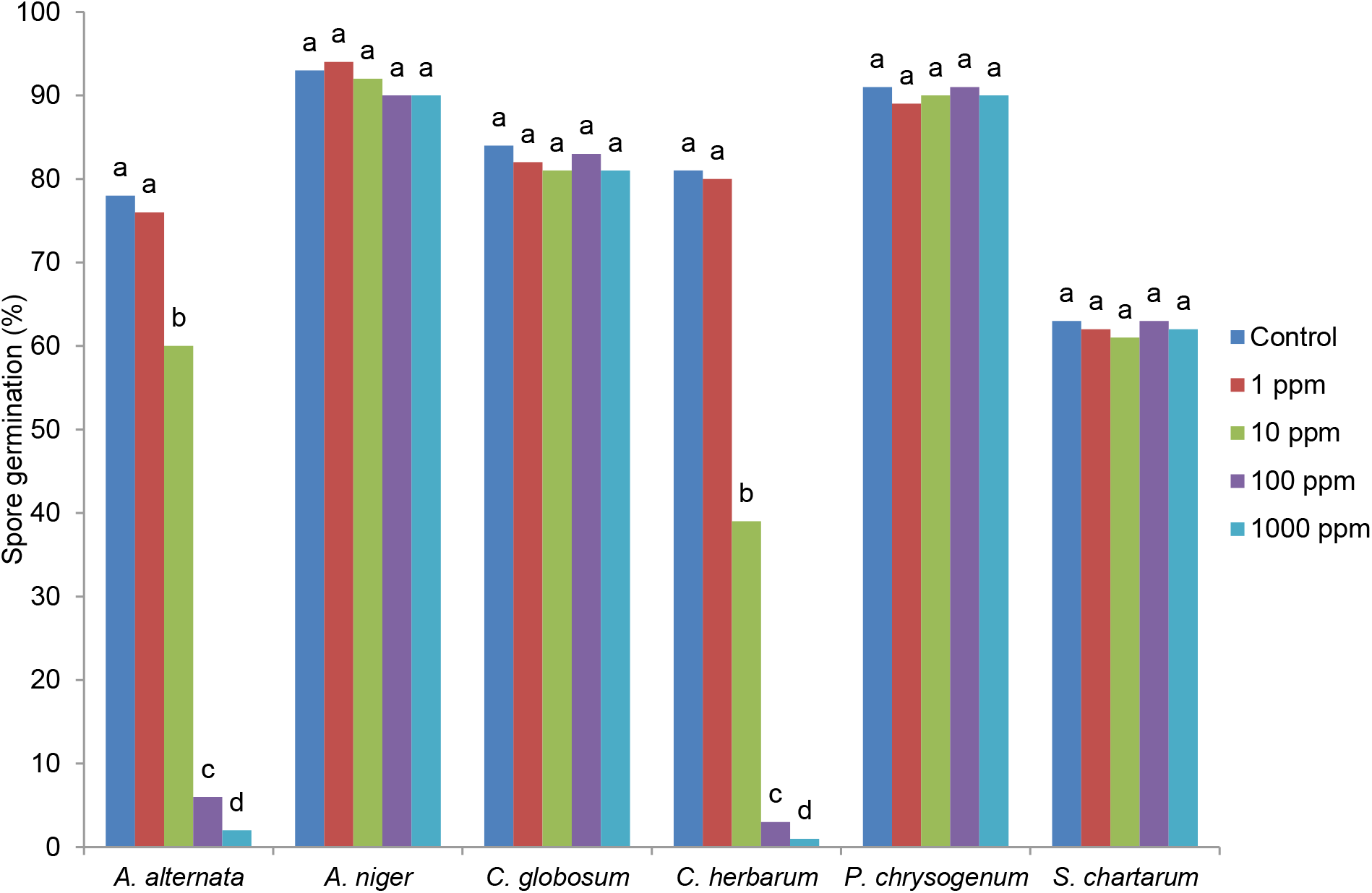
Effect of trichodermin on the spore germination of six fungal species. Values are the means of 20 replicates. Means followed by the same letters in bars (a,b,c, and so on) for a particular fungal species against trichodermin are not signicantly (P=0.05) different from each other by Scheffe’s test for multiple comparison.

**FIG. 2.**
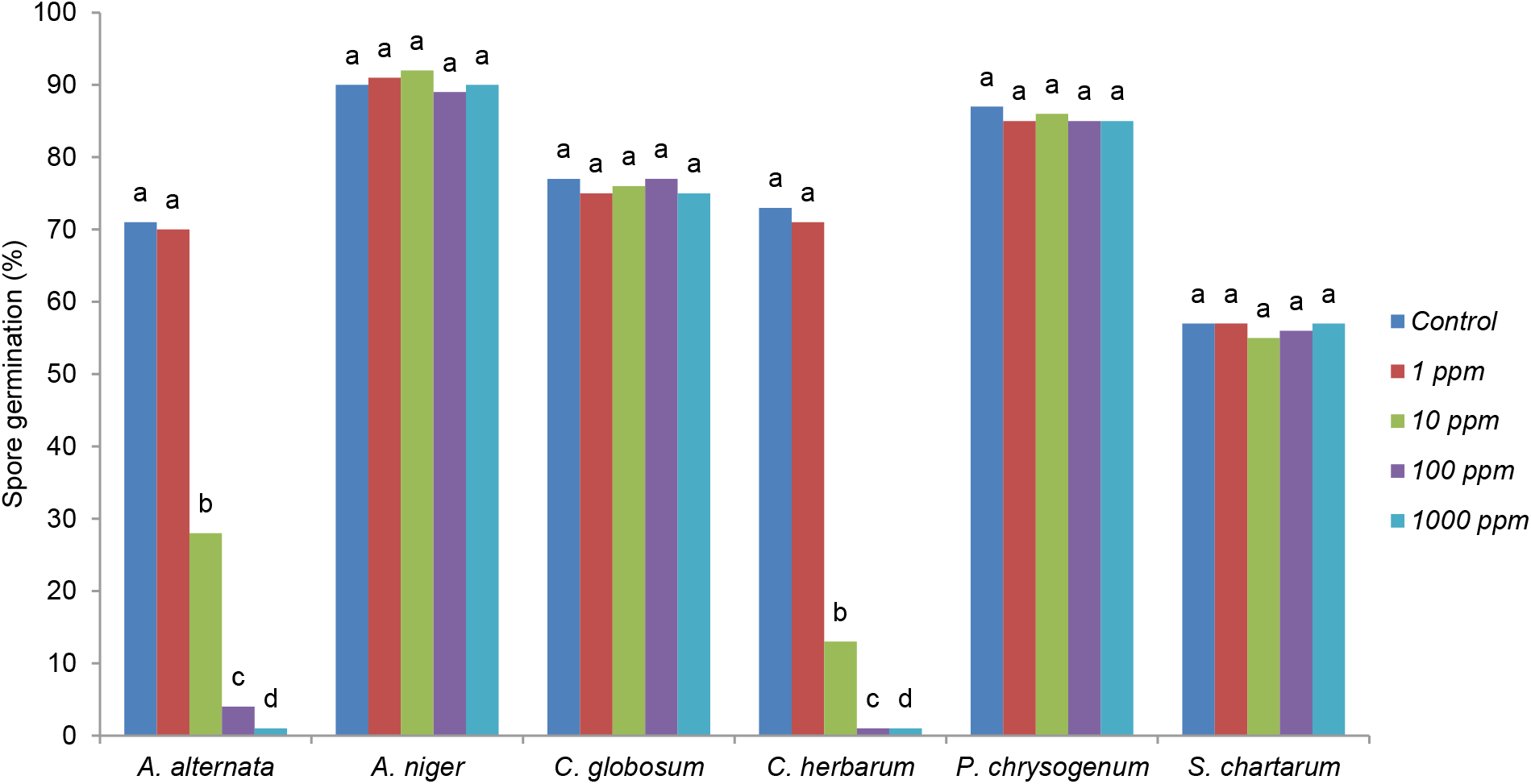
Effect of trichodermol on the spore germination of six fungal species. Values are the means of 20 replicates. Means followed by the same letters in bars (a,b,c, and so on) for a particular fungal species against trichodermol are not signicantly (P=0.05) different from each other by Scheffe’s test for multiple comparison.

#### Effect of trichodermin and trichdermol on the *in vitro* mycelial growth of six species of fungi

Both trichodermin and trichodermol at 1 ppm had no affect on the *in vitro* growth of *A. alternata* and *C. herbarum* (Table 4). *In vitro* growth of *A. alternata* and *C. herbarum* was significantly reduced when treated with trichodermin and trichodermol at 10, 100, and 1000 ppm (Table 4). Both these compounds had no effect on *in vitro* growth of *A. niger, C. globosum, P. chrysogenum* and *S. chartarum* at 1, 10, 100, and 1000 ppm.

**TABLE 4.** Effect of trichodermin and trichodermol on the *in vitro* mycelial growth of six fungal species on different nutrient media

| Fungal species | Nutrient media | % Reduction of growth from control |  |  |  |  |  |  |  |
| --- | --- | --- | --- | --- | --- | --- | --- | --- | --- |
|  |  | Trichodermin (ppm) |  |  |  | Trichodermol (ppm) |  |  |  |
|  |  | 1 | 10 | 100 | 1000 | 1 | 10 | 100 | 1000 |
| *A. alternata* | MEA | 0.0d | 13.0b | 100a | 100a | 0.0d | 5.5c | 100a | 100a |
|  | PDA | 0.0d | 22.5b | 100a | 100a | 0.0d | 9.0c | 100a | 100a |
|  | CA | 0.0d | 19.5b | 100a | 100a | 0.0d | 4.0c | 100a | 100a |
| *A. niger* | MEA | 0.0a | 0.0a | 0.0a | 0.0a | 0.0a | 0.0a | 0.0a | 0.0a |
|  | PDA | 0.0a | 0.0a | 0.0a | 0.0a | 0.0a | 0.0a | 0.0a | 0.0a |
|  | CA | 0.0a | 0.0a | 0.0a | 0.0a | 0.0a | 0.0a | 0.0a | 0.0a |
| *C. globosum* | MEA | 0.0a | 0.0a | 0.0a | 0.0a | 0.0a | 0.0a | 0.0a | 0.0a |
|  | PDA | 0.0a | 0.0a | 0.0a | 0.0a | 0.0a | 0.0a | 0.0a | 0.0a |
|  | CA | 0.0a | 0.0a | 0.0a | 0.0a | 0.0a | 0.0a | 0.0a | 0.0a |
| *C. herbarum* | MEA | 0.0d | 19.0b | 100a | 100a | 0.0d | 6.5c | 100a | 100a |
|  | PDA | 0.0d | 10.5b | 100a | 100a | 0.0d | 3.0c | 100a | 100a |
|  | CA | 0.0d | 13.5b | 100a | 100a | 0.0d | 2.0c | 100a | 100a |
| *P. chrysogenum* | MEA | 0.0a | 0.0a | 0.0a | 0.0a | 0.0a | 0.0a | 0.0a | 0.0a |
|  | PDA | 0.0a | 0.0a | 0.0a | 0.0a | 0.0a | 0.0a | 0.0a | 0.0a |
|  | CA | 0.0a | 0.0a | 0.0a | 0.0a | 0.0a | 0.0a | 0.0a | 0.0a |
| *S. chartarum* | MEA | 0.0a | 0.0a | 0.0a | 0.0a | 0.0a | 0.0a | 0.0a | 0.0a |
|  | PDA | 0.0a | 0.0a | 0.0a | 0.0a | 0.0a | 0.0a | 0.0a | 0.0a |
|  | CA | 0.0a | 0.0a | 0.0a | 0.0a | 0.0a | 0.0a | 0.0a | 0.0a |

#### Effect of *S. chartarum* and five fungal species on colonization in building materials

All the six species of fungi grew well in drywall, ceiling tile, and oak wood chips (Table 5). Both *A. alternata* and *C. herbarum* failed to grow in these substrate when grown together with *S. chartarum* (Table 5). The growth of A. *niger, C. globosum,* and *P. chrysogenum* was not inhibited when grown together with *S. chartarum* in these substrate (Table 5).

**TABLE 5.**
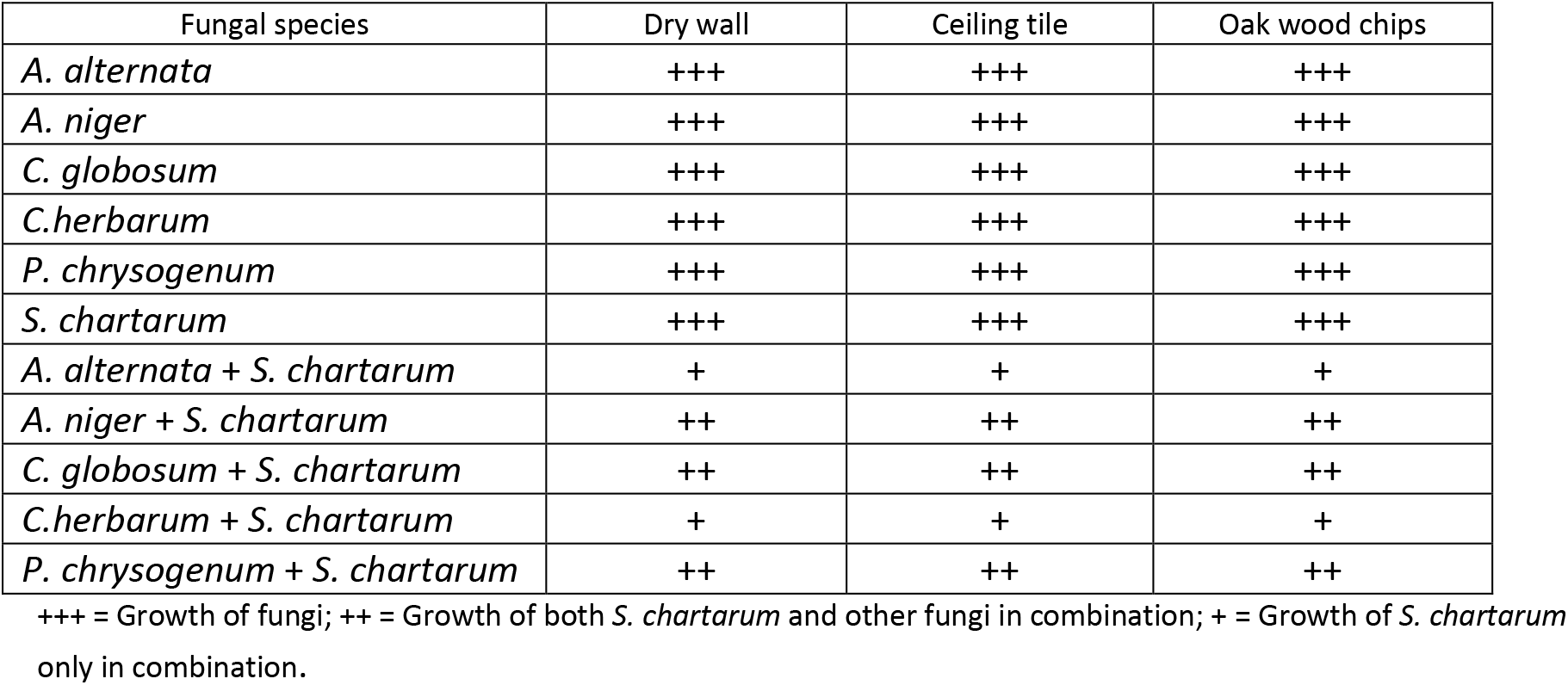
Growth of six fungal species and interaction of *Stachybotrys chartarum* with other fungal species in the building materials

## DISCUSSION

The present study was designed to investigate the synergistic interactions between *S. chartarum* and other commonly occurring indoor fungal species in water damaged building materials. Modern building materials, once moistened, may provide rich ecological niches for various fungi. It has long been postulated that damp or homes with high humidity have a musty smell or have obvious fungal growth. The occupants of these homes have increase in subjective complains including upper respiratory, asthma, gastrointestinal, and other illness (10,22,29,30). During this study, we have found that the most common fungal species in the water damaged building materials in the southern California were species of *Alternaria, Aspergillus, Chaetomium, Cladosporium, Penicillium* and *Stachybotrys*. Interestingly, it was observed that *A. alternata* and *C. herbarum* were always absent when *S. chartarum* was growing on the same building materials. On the other hand, colonies of *A. niger, C. globosum*, and *P. chrysogenum* were often associated with *S. chartarum*.

*In vitro* inhibition of fungi has been attributed through parasitism or surface contact followed by penetration and formation of intercellular hyphae within the host hyphae, or production of antifungal compounds by antagonist fungus (11). In this study, *S. chartarum* was an effective antagonist against *A. alternata* and *C. herbarum* owing to the production of antifungal substances that were excreted in the culture media. *Stachybotrys chartarum* was capable of inhibiting growth of both *A. alternata* and *C. herbarum.* Although *S. chartarum* did not parasitize or penetrate the hyphae of *A. alternata* and *C. herbarum,* it appears to kill these two fungi through the production of metabolites. The cytoplasm of *A. alternata* and *C. herbarum* were coagulated and disintegrated when exposed to *S. chartarum*. In this study, mycelial growth of *A. alternata* and *C. herbarum* was also inhibited in both agar and liquid culture by *S. chartarum*. The crude extract of *S. chartarum* also strongly reduced the growth of *A. alternata* and *C. herbarum.* These results indicate that inhibition of *A. alternata* and *C. herbarum* by *S. chartarum* is not hyphal parasitism rather it is a chemical in nature.

All these six fungal species described above are known to produce mycotoxin. Mycotoxins are the secondary metabolites of fungi that represent a chemically diverge group of organic, non-volatile, and low molecular weight compounds. Mycotoxins are usually produced when conditions favor fungal growth such as moisture, pH, growth medium, and temperature. Among these fungi, *S. chartarum* is considered to be one of the most toxic fungi and produce cytotoxic compound known as trichothecene. Trichothecenes are secondary metabolites produced by species of *Stachybotrys* as well as other fungi that are harmful to human and animal health causing wide range of diseases (9,14,26). There are over 150 trichothecenes and trichothecenes derivatives have been isolated and characterized (14). They are all non-volatile, low molecular weight sesquiterpene epoxides and share a tricyclic nucleus and usually contain an epoxide at C-12 and C-13, which are essential for toxicity (12). The trichothecene skeleton is chemically stable and not degraded by heat or neutral or acidic pH (13). In this study, two trichothecene, trichodermin and trichodermol produced by species of *Stachybotrys* showed significantly inhibitory effect on *A. alternata* and *C. herbarum* at 10, 100, and 1000 ppm. Both spore germination and mycelial growth of these fungi were significantly inhibited by these compounds. Interestingly, *A. niger, C. globosum, P. chrysogenum*, and *S. chartarum* were not affected by trichodermin and trichodermol up to 1000 ppm indicating that sensitivity of these trichothecenes varied considerably amongst different fungal species.

The antifungal activity of *S. chartarum* was also confirmed on drywall, ceiling tile, and oak wood chips where it prevented the growth of *A. alternata* and *C. herbarum* in these substrates. For antibiotic metabolite production to occur in these substrate, certain conditions for secondary metabolism would have to met. Secondary metabolite formation usually occurs after a period of hyphal growth has taken place, thus establishing a certain hyphal mass, age, or growth rate (8). It is interesting to note that growth of *S. chartarum* was not affected by its own metabolite. Both *A. alternata* and *C. herbarum* colonized abundantly in building materials in the same homes when *S. chartarum* was absent, however, they failed to colonized in the same building materials when *S. chartarum* was present.

This study shows that there exist an associated mycobiota on the damp building materials based on interactions between these fungi. The first association of fungi was between *A. alternata, A. niger, C. globosum, C. herbarum* and *P. chrysogenum*. Although all the fungi produce mycotoxins these fungi have found their niche on damp or wet building materials without inhibiting growth of each other. A second strong association was seen between mycotoxin producing fungi *A. niger, C. globosum, P. chrysogenum*, and *S. chartarum* and they seem to coexist together on damp or wet building materials.

From this study, distinct interaction patterns between *S. chartarum* against *A. alternata* and *C. herbarum* was identified and determined by the antifungal compounds production by *S. chartarum.* Trichodermin and trichodermol were toxic to both *A. alternata* and *C. herbarum* at concentrations 10 ppm and above but had no affect on *A. niger, C. globosum,* and *P. chrysogenum* up to 1000 ppm. This may explains why *A. alternata* and *C. herbarum* were absent in the same building materials when colonized by *S. chartarum* but the growth *A. niger, C. globosum,* and *P. chrysogenum* was not affected. This study demonstrates that synergistic interaction between different fungi play an important role in the colonization of damp or wet building materials.

## ACKNOWLEDGMENTS

This study was supported by Pasteur Laboratory’s R & D programs. I would like to thank staff members of the laboratory for their valuable advice and useful suggestions on the manuscripts. The helpful field works by the team members of the Safegurad EnviroGroup are gratefully acknowledged.

